# Targeting wild type NTRK decreases brain metastases of lung cancers non-driven by NTRK fusions

**DOI:** 10.64898/2026.03.18.711213

**Authors:** Maria J. Contreras-Zárate, Jenny A. Jaramillo-Gómez, R. Alejandro Marquez-Ortiz, Trinh C. Pham, Stella Koliavas, D. Ryan Ormond, Andre C. Navarro, Raphael A Nemenoff, D. Ross Camidge, Diana M. Cittelly

## Abstract

The central nervous system (CNS) is a common site of metastatic spread for both non-small cell and small cell lung cancer, yet the therapeutic strategies to prevent and decrease lung cancer brain metastases remain limited. Tyrosine kinase inhibitors have shown promising results in increasing the overall response in brain metastases, owing to their brain penetrance and increased effectiveness; however, their use is limited to the small group of tumors carrying specific oncogenic drivers. Among these, inhibitors with activity against neurotrophic tyrosine receptor kinases (NTRKs) are showing promising effects in reducing CNS metastases in cancers driven by gene rearrangements of these drugs’ targets. However, wild-type NTRKs are susceptible to activation by their canonical ligands, which are expressed throughout the brain metastatic niche and can, in a paracrine manner, activate NTRK function in cancer cells. Here we show that NTRKs are expressed in primary tumors, brain metastases, and lung cancer cells with various driver mutations expressing wild-type NTRK2 (WT-TrkB). We demonstrate that WT-TrkB activates downstream signaling and proliferation in response to exogenous BDNF and conditioned media from reactive astrocytes known to secrete BDNF in the brain niche. Importantly, the FDA-approved NTRK inhibitor entrectinib blocked BDNF and astrocyte-induced survival pathways in multiple lung cancer cell lines, decreased their proliferation *in vitro*, and effectively prevented brain metastatic colonization and progression *in vivo* without significant effects on extracranial disease. Thus, these studies suggest that brain-dependent activation of NTRK is critical for brain metastases of WT-NTRK+ lung cancers, and therefore, NTRK inhibitors can be used to target non-fusion NTRK function to prevent or decrease brain metastases.

**SIGNIFICANCE:** These studies demonstrate that NTRK wild-type receptors are important drivers of brain metastatic colonization and progression in different subtypes of lung cancer, independent of their driver alterations. Thus, they provide rationale to expand the use of FDA-approved NTRK inhibitors with brain penetrance for the prevention of CNS metastases.

## INTRODUCTION

Brain metastases (BM) are a common site of metastatic spread for both non-small cell and small cell lung cancer, affecting 20-56% of patients with advanced lung cancer^1^. The increase in lung cancer BM is likely the result of better neuroimaging capabilities and longer patient survival driven by effective new cancer treatments, which provide greater opportunities for the disease to spread to the brain. Treatment strategies for BMs, once dominated by local methods like surgery, whole-brain radiotherapy (WBRT), and stereotactic radiosurgery (SRT), have been reshaped by the emergence of more effective systemic options including targeted therapies, immune checkpoint inhibitors and antibody drug conjugates^2^. Yet the mortality associated with lung cancer BM progression remains high and strategies to prevent or decrease CNS involvement remain an urgent unmet need.

Among emerging therapeutic strategies for lung cancer BM, tyrosine kinase inhibitors have shown promising intracranial activity, in part due to their ability to penetrate the brain and effectively inhibit tumors driven by their specific oncogenic alterations^3^. Importantly, inhibitors targeting the NTRK family, which includes NTRK1, NTRK2 and NTRK3, encoding TrkA, TrkB and TrkC, respectively, either as direct targets (e.g Larotrectinib) or secondary targets (e.g. entrectinib, reprotectinib, lorlatinib which also inhibit ALK and ROS1), offer new opportunities for tumors expressing wild type NTRK receptors^4,5^. These receptors can be activated by their cognate ligands, which are highly abundant in the CNS^4,6^, supporting tumor growth within the brain microenvironment^7^. However, the degree to which pharmacological NTRK inhibition can affect lung cancer brain metastases in tumors lacking NTRK gene rearrangements or other oncogenic targets of these drugs remains unclear.

Among NTRKs, wild-type TrkB expression has been shown to promote lung adenocarcinoma metastases^8^ and TrkB expression has been associated with worse survival in a panel of non-small cell lung cancer (NSCLC) samples containing squamous cell carcinomas, adenocarcinomas, and large-cell neuroendocrine carcinomas^9,10^. TrkB in its native form binds with high affinity to its cognate ligand brain-derived neurotrophic factor (BDNF) and to a less extent to Neurotrophin-4 (NT-4). BDNF/TrkB pro-tumorigenic function is well recognized in tumors with propensity to colonize the CNS (breast, lung, colorectal), playing critical roles in the ability of disseminated cancer cells to resist anoikis, survive and invade at metastatic sites^8,11–15^. BDNF is expressed at a high level in certain regions in the brain, particularly hippocampus, cerebral cortex, basal forebrain, and striatum, and its expression is strongly regulated by neuronal activity^16,17^. Astrocytes also secrete BDNF in response to multiple stimuli^18–21^ and we have shown that breast cancer cells expressing wildtype TrkB activate downstream signaling in response to astrocytic-BDNF, and that the BDNF/TrkB inhibitor ANA-12 prevents brain metastatic colonization of TrkB+ BC cells^7^. Here, we demonstrate that TrkB, and to a lesser extent TrkA, are expressed in multiple lung cancer cells without NTRK rearrangements. WT-TrkB is activated in response to exogenous BDNF, astrocytic BDNF and the brain niche, and can be blocked to different extents with pan TRK inhibitors. We show that astrocytes in the brain niche support growth of lung cancer cells, and entrectinib, a pan TRK inhibitor with high brain penetrance^22^, can prevent and decrease progression of non-NTRK rearranged lung cancer brain metastases in pre-clinical models. Thus, these studies provide preclinical data to extend the use of NTRK inhibitors to target brain-specific survival pathways.

## METHODS

### Cell Lines

Human NSCLC cell lines LU65 (RRID:CVCL_1392), H358 (RRID:CVCL_1559), both harboring heterozygous KRAS^G12C^ mutations, and their GFP□/Luciferase□ derivatives were cultured in RPMI-1640 supplemented with 10% FBS and 1% penicillin–streptomycin (complete media). Murine NSCLC cells CMT167 (RRID:CVCL_2405) and their Luciferase□ derivatives were cultured in DMEM supplemented with 10% FBS, 1% penicillin–streptomycin, and 20 mM HEPES. Brain-tropic NSCLC lines PC9Br3 (EGFR Δexon19 mutation) and H2030Br3 (KRAS^G12C^) were cultured as described previously^23^. All experiments used cells within five passages from thawing. Mycoplasma testing was performed every three months using MycoAlert™ PLUS (Lonza).

### Cell proliferation assays

NSCLC cells (2,000 per well) were seeded in 96□well plates in 100 µL of starvation medium and incubated overnight. The starvation medium was composed of phenol red–free DMEM (4.5 g/L glucose) supplemented with penicillin/streptomycin (P/S), non□essential amino acids, sodium pyruvate, 0.1% BSA, and 5% charcoal□stripped FBS (cs□FBS). Next day, 50 µL of medium was removed and replaced with 50 µL of starvation medium containing either drugs (2X, 2 µM) or vehicle (DMSO), resulting in a final concentration of 1 µM. After 2 hours, treatments were applied with either BDNF (final concentration 50 ng/mL) or astrocyte□conditioned medium (Ast□CM; 20X stock diluted to a final concentration of 10X). Cells were imaged over time using the IncuCyte Live Cell Imaging system (Essen Bioscience). Cell confluence per well was quantified from 4 fields per well, with at least 5 replicates per treatment.

### Astrocyte co-cultures and conditioned media

Primary neonatal astrocytes were isolated from P0–P2 CD1 mouse pups and seeded in T175 flasks and cultured as previously described^24^. Astrocytes were cultured in regular medium (DMEM + 10% FBS + 1% P/S) until 100% confluence was reached, and microglia free cultures were used for co-culture experiments or obtaining conditioned media (mAst-CM). For *co-culture experiments*, astrocytes were seeded in 12-well plates, grown to 100% confluence and then cancer cells (10,000 cells/well) were seeded on top in 1 mL of starvation medium (DMEM supplemented with 2% CS□FBS) and incubated overnight. Treatments were applied at 2X concentration in 1 mL of serum□free starvation medium. Total Green Object Area (µm²/Image) was quantified from 16 fields per well, with 4 replicates per treatment. For obtaining *conditioned media*, T175 flasks at 100% confluence were rinsed with PBS and cultured on 25 mL of starvation medium for 72 hours. The conditioned media was collected, passed through a 0.2 µm filter, and concentrated 10X using 3kD cutoff Amicon Ultra centrifugal filters (15 mL, UFC900324). Freshly prepared Ast□CM was used in all experiments.

### Human Clinical Samples (IHC)

Archival paraffin-embedded lung cancer brain metastasis samples were obtained from the University of Colorado Department of Pathology Biorepository under COMIRB protocol 15-1461. Formalin-fixed, paraffin-embedded sections were deparaffinized, subjected to heat-mediated antigen retrieval, and incubated with anti-TrkB antibody (Proteintech) overnight, followed by ImmPRESS™ anti-rabbit IgG secondary antibody (Vector Laboratories) for 30 min at room temperature. Detection was performed using DAB (DAKO kit; 1 drop [20 µL] per 1 mL substrate buffer). Sections were counterstained with Vintage hematoxylin (1:1 dilution) for 1 min, dehydrated, cleared, and mounted with Permount™.

### Western Blot

Cells were lysed at 4□°C for 5□min in 2× RIPA buffer containing protease (Roche #04693159002) and phosphatase inhibitors (Roche #04906837001), followed by four 1s sonication pulses at 20% amplitude. Protein concentration was measured using the Bio-Rad DC Protein Assay Kit II (#5000112). Between 20–40□µg of protein were resolved at 100□V on 8–10% SDS-PAGE or 4–15% precast gels (Bio-Rad #4561086). Proteins were transferred to PVDF membranes (Immobilon-FL 0.45□µm, Millipore #IPFL00010 for >40□kDa; Hybond 0.2□µm, Amersham #10600022) and blocked with 3% BSA in TTBS for 1□h at room temperature before overnight incubation with specific primary antibodies (Supplementary Table 1).

### Animal Experiments

All procedures were approved by the University of Colorado IACUC. Investigators were blinded to treatment groups. NSG mice (8–12 weeks old, male or female) received intracardiac injections of 200,000 H2030Br3 GFPLuc□ cells. *Preventive Study:* Ovariectomized (OVX) female NSG mice supplemented with a 1 mg estradiol (E2) pellet were randomized to vehicle or Entrectinib (ENT; 60 mg/kg, BID, PO) beginning three days before intracardiac injection and continued for four weeks. *Therapeutic Study:* NSG male and female mice were injected intracardially as above and head IVIS signal was monitored every 10 days. Mice with head signal ≥2× baseline 3 weeks after injection were randomized to receive vehicle or ENT (60 mg/kg, BID, PO) for 14 days. In all studies metastatic burden was monitored weekly via IVIS.

### In Vivo Luminescence (IVIS)

Mice received 200 µL of D-Luciferin (GoldBio LUCK-1G; 15 mg/mL) via subcutaneous injection. Head and extracranial metastatic burden were quantified using *Living Image®* software (v2.60.1). At euthanasia, brains were incubated in 0.15 mg/mL Luciferin in PBS for 10 min and imaged ex vivo.

### Histological Quantification

Micro- (<300 µm) and macro-metastases (>300 µm) were quantified as described ^25^. Briefly, six H&E-stained serial sections, 300 µm apart in a sagittal plane through one hemisphere, were analyzed at ×4 magnification using an ocular grid. Median counts per mouse were recorded.

### Immunofluorescence analysis

Fresh OCT-embedded hemisphere brains were sectioned at 20 µm thickness and stored at −80 °C until use. Slides were thawed at room temperature and fixed in cold acetone for 5 min, followed by rinsing in TBST. Sections were blocked with 10% normal donkey serum (NDS) for 30 min. Primary antibodies were incubated overnight (∼16 h) under the conditions specified in **Supplementary Table 1**. Secondary antibodies were incubated for 1 h at room temperature. Slides were mounted using Fluoromount-G containing DAPI (Invitrogen 00-4959-52). Images were acquired using an Olympus APX100 with a 10X objective for whole-brain images and a 40X objective for regions used in analysis. Image quantification was performed using ImageJ.

#### Image Analysis and Quantification

Fluorescence images were analyzed using a custom macro in ImageJ. the GFP channel was used to generate a segmentation mask. Thresholding of the GFP image was manually adjusted, after which the image was converted to a binary mask and refined using hole filling, dilation, and erosion. The resulting mask was converted to a region of interest (ROI) and applied to the corresponding RFP image to measure mean RFP fluorescence intensity and area within the GFP-positive region. Measurements from all samples were automatically compiled into a single CSV file for statistical analysis.

### Publicly available data analysis

Gene expression data for lung adenocarcinoma (Project ID: TCGA-LUAD) were obtained from The Cancer Genome Atlas (TCGA) using the TCGA biolinks R package^26^. Gene Expression Quantification counts were downloaded as normalized fragments per kilobase of transcript per million mapped reads (FPKM). Expression profiles for NTRK1, NTRK2, and NTRK3 were extracted specifically from samples categorized as tumor tissue.

The exported dataset was imported into GraphPad Prism version 10.6.1 for statistical analysis. Differences in transcriptional levels among the three NTRK genes were assessed using ordinary one-way analysis of variance (ANOVA) followed by Tukey’s multiple comparisons test to determine pairwise significance.

Spatial transcriptomic data were obtained from the publicly available GSE200563 dataset in the NCBI GEO^27^. Raw Digital Spatial Profiling (DSP) counts were processed in R to correct for technical variability across Areas of Interest (AOIs) using 75th-percentile (Q3) normalization. Expression values for NTRK1 and NTRK2 were extracted and paired by patient identifier to enable direct comparison between primary tumors (PT) and their matched BM in NSCLC patients (n = 23). Statistical analyses were performed using the Wilcoxon signed-rank test to evaluate paired differences in gene expression between PT and BM samples.

### Statistical Analysis

Statistical tests and sample sizes are indicated in figure legends. Analyses were performed using *GraphPad Prism* v10.6.1. Data normality was assessed by Shapiro–Wilk and Kolmogorov–Smirnov tests. Parametric data were analyzed by one-way ANOVA or unpaired t-tests; non-parametric data by Kruskal–Wallis or Mann–Whitney U tests with appropriate multiple-comparison corrections. P ≤ 0.05 was considered significant.

## RESULTS

### TrkB is expressed in lung cancer cells and is activated by exogenous BDNF

We first sought to confirm the extent to which NTRK are expressed in lung cancer patients and their brain metastases. Analysis of publicly available datasets (TCGA-LUAD) showed NTRK1, NTRK2 and NTRK3 mRNA expression in primary human lung adenocarcinoma, with higher expression of NTRK2 mRNA (**Fig 1A**). Comparison of NTRK2 mRNA levels by spatial transcriptomic analysis (GSE200563) in a cohort of lung primary tumors and their matched brain metastases^27^ shows similar expression of NTRK1, NTRK2 and NRTK3 in both primary and brain metastases, suggesting NTRK2 expression is a property intrinsic to the primary tumors (**Fig 1B**). TrkB protein expression was confirmed by immunohistochemistry in a small cohort of lung cancer brain metastases (n=9) without NTRK driver mutations (**Fig 1C**). We confirmed that WT-TrkB is more prevalent than TrkA and TrkC across a panel of cell lines, and detected variable expression of full-length TrkB (110 to 145 KD, consistent with differential glycosylation)^28^ in human (LU65, PC9Br3, H358 and H2030Br) and murine (CMT167) NSCLC cells, all of which lack NTRK rearrangements (**Fig. 1D**). As previously reported, BDNF is expressed in some lung cancer cell lines as well as in human and murine astrocytes in the normal brain^29,30^, and in reactive astrocytes surrounding lung cancer brain metastases^7^ (**Fig. 1E, F**) suggesting that both autocrine and paracrine mechanisms may activate cancer cell NTRK signaling. To assess whether WT-TrkB is responsive to exogenous BDNF stimulation, we conducted a time-course analysis of TrkB activation and downstream signaling in TrkB+ lung cancer cell lines. BDNF induced a robust activation of TrkB downstream signaling, characterized by increased phosphorylation of AKT and ERK within 5-10 min (**Fig. 1G**). Notably, TrkB and downstream AKT phosphorylation were observed in the absence of exogenous BDNF in some cell lines, especially PC9Br3, likely reflecting their high levels of endogenous BDNF expression (**Fig 1E**).

**Fig. 1.**
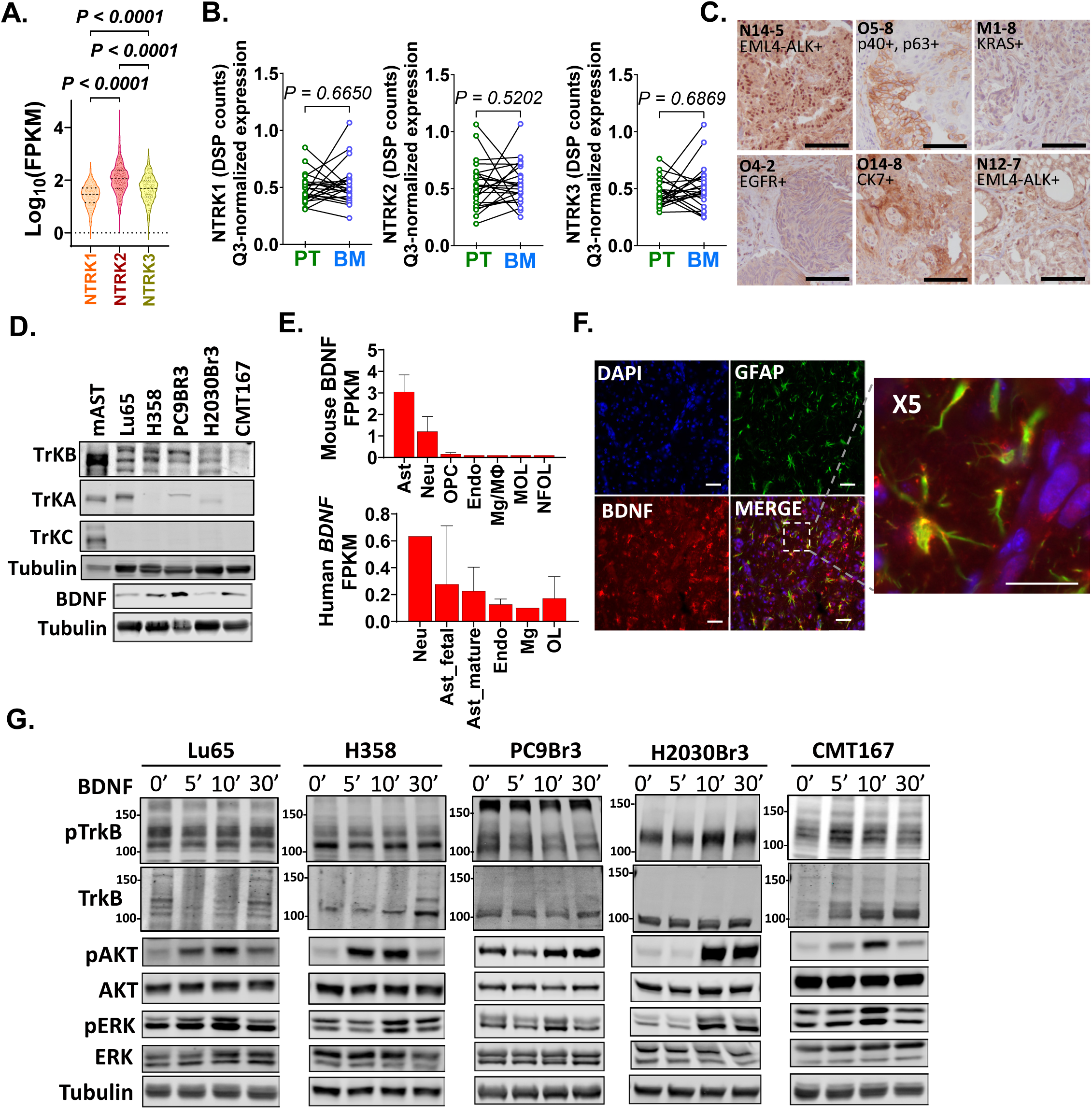
TrkB is expressed in lung cancer cells and activated by exogenous BDNF. **(A)** Gene expression levels of NTRK1, NTRK2, and NTRK3 in lung adenocarcinoma (LUAD) tumor samples from The Cancer Genome Atlas (TCGA). Normalized gene counts (FPKM) were obtained using the TCGAbiolinks R package and analyzed in GraphPad Prism 10.6.1. One-way ANOVA with Tukey’s multiple comparisons **(B)** NTRK1 and NTRK2 expression between primary tumors (PT) and matched brain metastases (BM) Q3-normalized values were paired by patient ID (n = 23). Paired comparisons were performed using the *Wilcoxon signed-rank test.* **(C)** Immunohistochemistry showing TrkB expression in a small cohort of NSCL-BM, scale bar = 100µm. **(D)**. Western blot showing expression of TrkB, TrkA, TrkC, and BDNF in NSCLC cell lines. Mouse astrocytes were used as a positive control of TrkA and TrkC. **(E)** Relative mRNA expression of mouse and human BDNF from Brain-RNAseq.org. **(F).** Immunofluorescence expression of BDNF in GFAP+ astrocytes surrounding lung cancer brain metastases in a xenograft mode. Scale bar 50 um **(G).** Western blots show the time course of BDNF-induced TrkB signaling in NSCLC cells serum-starved overnight and stimulated with 50 ng/mL BDNF. Tubulin served as a loading control in all blots. 150 and 100 KD molecular weight markers show full length TRKB expression (110 to 140 KD depending on glycosylation levels).

### Entrectinib blocks BDNF induced TrkB activation and proliferation of TrkB+ NSCLC cells without NTRK driving mutations

To evaluate whether tyrosine kinase inhibitors targeting TrkB (Entrectinib, LOXO 101, Cyclotraxin, and ANA12) could block BDNF-induced TrkB signaling, H358, H2030Br3, and CMT167 cells were serum-starved overnight and pretreated with 1 µM TKI for 2 hours, prior to stimulation with exogenous BDNF for 10 min. Among the inhibitors tested, entrectinib effectively reduced BDNF-induced AKT and ERK activation in H358, H2030Br3, and CMT167 cells, but not in Lu65 or PC9Br3 cells (**Fig 2A, Sup. Fig 1A**). In triple negative breast cancer models (4T1Br5 and E0771) where we had previously demonstrated that BDNF/TrkB signaling is important for proliferation and migration functions, entrectinib was also able to reduce BDNF-induced activation of AKT and ERK (**Sup. Fig 1A**), indicating the TRK-B blocking function of entrectinib encompasses multiple CNS-prone primary tumors.

**Fig. 2.**
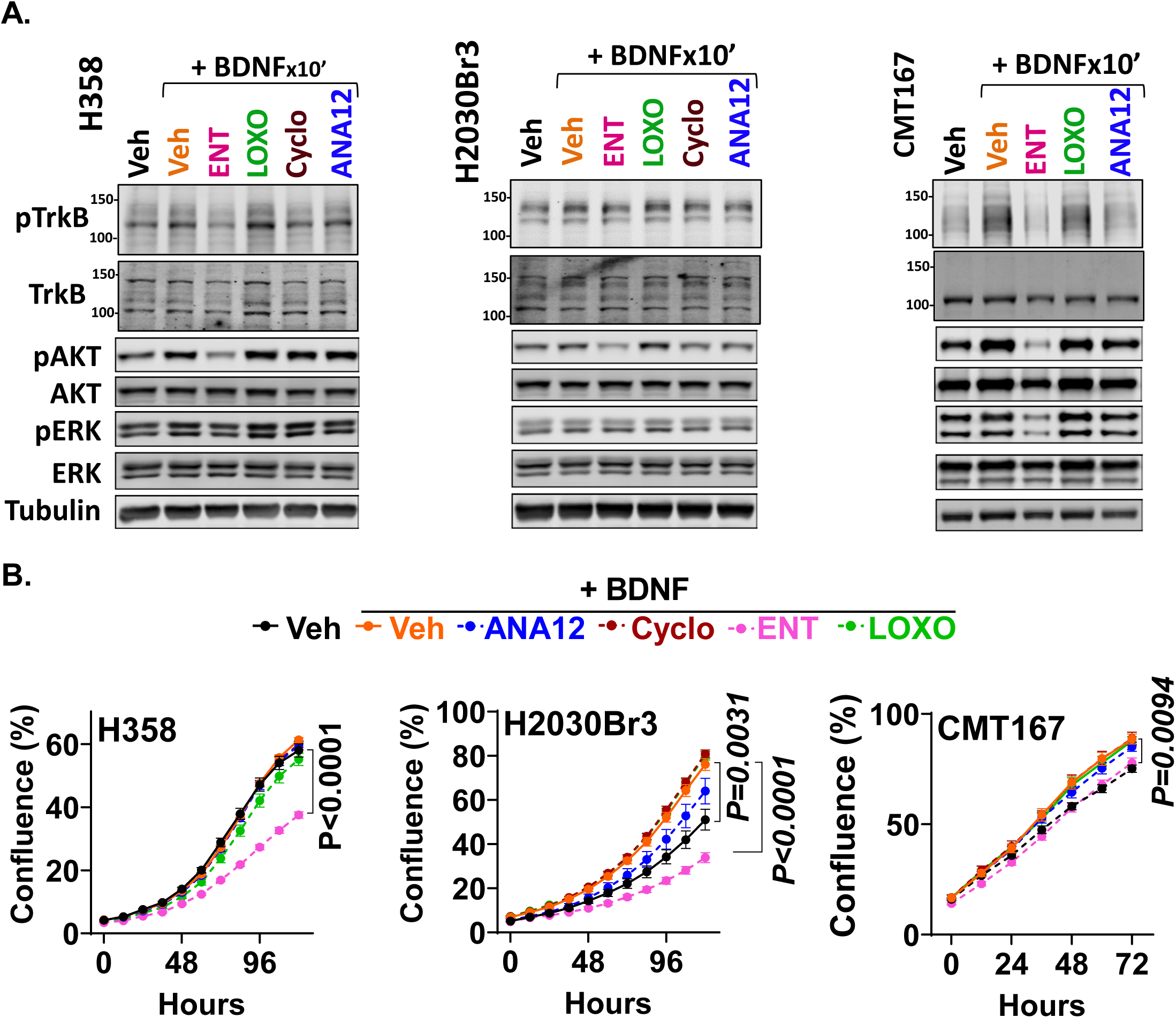
Entrectinib blocks TRKB activation and proliferation of NSCLC TRKB⁺ cells lacking NTRK driving mutations. **(A)** Western blot of NSCLC cells plated in phenol-red–free media with 2.5% CS-FBS and pretreated for 2 hours with DMSO (Veh), Entrectinib (ENT), LOXO101 (LOXO), Cyclotraxin (Cyclo), or ANA12 (all drugs at 1 μM), followed by stimulation with 50 ng/mL BDNF for 10 minutes. Tubulin served as a loading control. **(B)** Proliferation measured as percent confluency (mean ± SEM) using Incucyte live-cell imaging (n = 5 per treatment). Cells were plated in phenol-red–free media with 2.5% CS-FBS and treated with vehicle, 50 ng/mL BDNF alone, or BDNF combined with 1 μM tyrosine kinase inhibitors. Data were analyzed by repeated-measures two-way ANOVA with post hoc multiple-comparison corrections.

To determine how BDNF/TrkB signaling impacts proliferation of lung cancer cells *in vitro*, cells were treated with BDNF alone or in combination with 1µM TKIs and cell proliferation was measured via live cell imaging. Exogenous BDNF significantly increased proliferation in cells with low intrinsic BDNF expression H2030Br3 and CMT167, but not in H358 cells. Consistent with the downstream inhibition of AKT in these cell lines, entrectinib significantly decreased their proliferation (**Fig 2B**). By contrast, while entrectinib did not alter BDNF-induced AKT and ERK activation in LU65 and PC9Br3 cells at the time-points tested (**Sup. Fig. 1B**), sustained entrectinib treatment inhibited proliferation in these cell lines (**Sup. Fig 1C**). Notably, entrectinib decreased proliferation of all cell lines (except for CMT167 cells) to levels below the vehicle control. Given that none of these cell lines express other entrectinib targets (ROS, ALK) (**Sup. Fig 1D**), this suggests that entrectinib can target proliferation through BDNF-independent NTRK activity (**Fig 2B**) and its consistent with its activity as a ATP-competitive kinase inhibitor able to shut down catalytic activity regardless of ligand availability, receptor conformation, or compensatory signaling feedback. ANA-12, a specific BDNF/TrkB competitive inhibitor was only effective in decreasing BDNF-induced proliferation in H2030Br cells, suggesting TrKB activation is an important driver of proliferation in these cells. Surprisingly, equivalent micromolar concentrations of other NTRK inhibitors (LOXO-101, Cyclotraxin-B) were unable to effectively decrease signaling or proliferation in most cell lines, suggesting entrectinib achieved robust pathway inhibition at the same dose.

### Entrectinib blocks proliferative signaling and cell proliferation of NSCLC cells in models mimicking interactions with the brain TME

We have reported that conditioned media from reactive astrocytes (Ast-CM) is a key source of growth factors, including, but not limited to, BDNF, and that they can, in a paracrine manner, activate pro-survival pathways and increase proliferation of brain metastatic breast cancer cells ^7,31^. To test how secreted factors from astrocytes impacted signaling and proliferation, lung cancer cells were serum-starved overnight, pretreated with 1µM TKI, and then treated with Ast-CM (10X) or control (1% CS-FBS media). Confirming prior findings in BC cells, Ast-CM strongly activated pro-survival AKT signaling pathways in H358, LU65 and PC9Br3 cells, moderately in H2030Br3, and did not impact AKT activation in CMT167 cells which showed strong levels of constitutive AKT activation **(Fig 3A, Sup. Fig 2A**). Since pretreatment with 1µM entrectinib but not 1µM LOXO-101 modestly reduced signaling induced by Ast-CM across multiple cell lines (despite significant decrease when cells where treated with BDNF alone (**Fig 2A**), we focused our next studies on defining the extent to which entrectinib could block or decrease activation of survival pathways in lung cancer cells in settings that mimic more complex interactions with the brain microenvironment. In H358 cells, entrectinib reduced Ast-CM-induced ERK activation but not AKT activation; while in H2030Br3 cells, entrectinib reduced endogenous but not Ast-CM induced AKT or ERK activation. By contrast, entrectinib effectively blocked both AKT and ERK activation in CMT167 and PC9Br3 cells alone or stimulated with Ast-CM (**Fig. 3A, Sup.2A**). Despite the differential effects of entrectinib on AKT and ERK activation among cell lines at the tested time points, sustained treatment with 1µM entrectinib significantly blocked cancer cell proliferation of vehicle-treated H358, H2030Br and Lu65 cells and reduced their growth in response to Ast-CM to levels similar to those of vehicle treated controls (**Fig. 3B, Sup.Fig. 2B**).

**Fig. 3.**
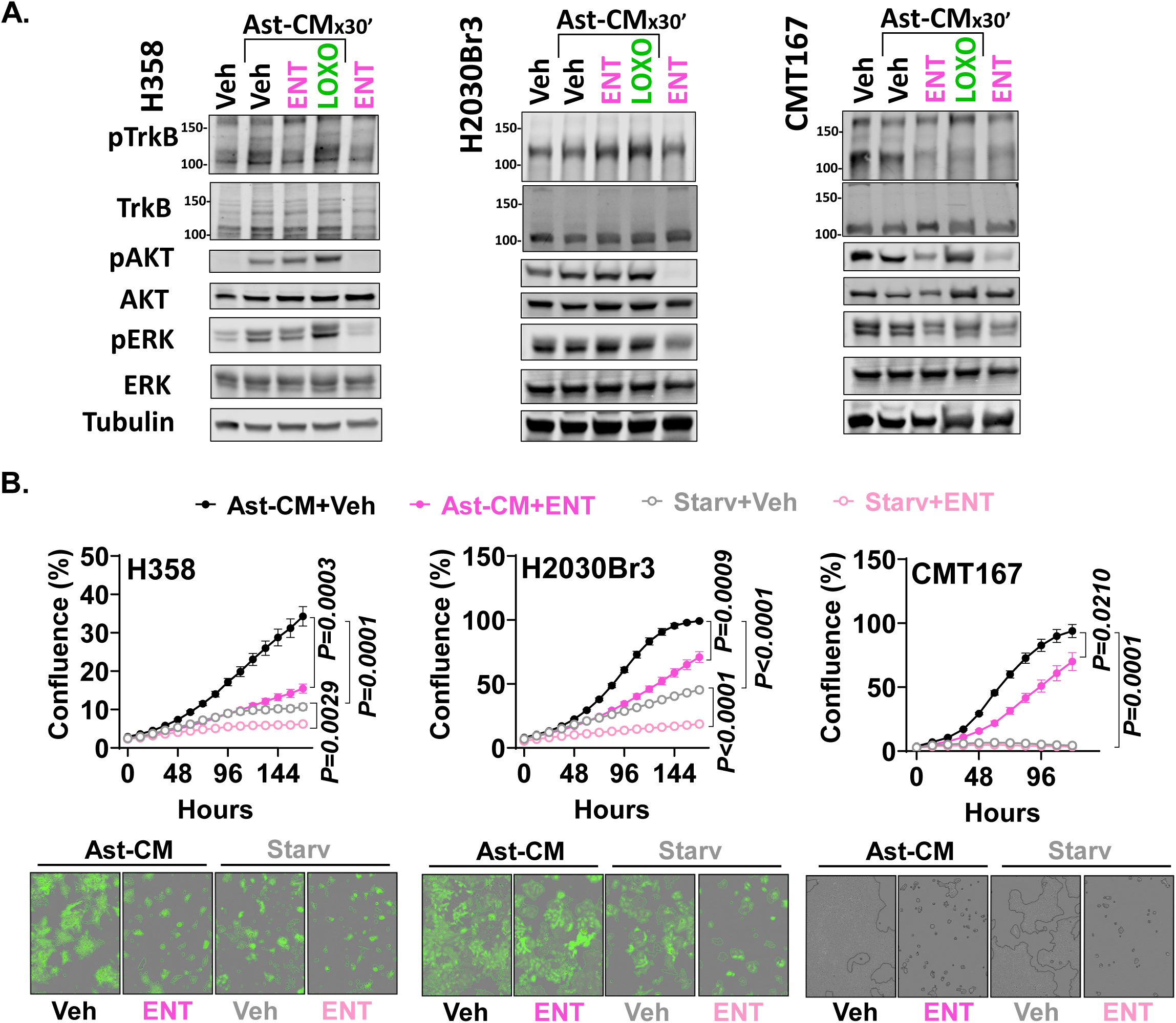
Entrectinib decreases AKT and ERK activation and proliferation induced by astrocyte-conditioned media (Ast-CM) in NSCLC cells. **(A)** Western blot of NSCLC cells serum-starved overnight, pretreated for 2 hours with 1 μM Entrectinib (ENT) or LOXO101 (LOXO), then stimulated with Ast-CM (10×) for 30 minutes. Tubulin served as a loading control. **(B)** Proliferation expressed as percent confluence (mean ± SEM) in NSCLC cells cultured in serum-starved media and treated with Ast-CM (10×) plus vehicle (DMSO) or ENT (1 μM). Data were analyzed by repeated-measures two-way ANOVA followed by Fisher’s LSD test (n = 5 wells per treatment).

CMT167 and PC9Br3 cells also showed decreased proliferation in response to entrectinib when cultured with Ast-CM, yet this growth was still significantly higher than vehicle treated cells, suggesting Ast-CM promotes additional growth in these cells through pathways that are not dependent on NTRKs.

Prior studies have shown that cell-cell contact with astrocytes can elicit pro-survival mechanisms in cancer cells different to those of secreted factors and can protect cancer cells from cytotoxic drugs^32,33^. To evaluate if entrectinib could decrease proliferation when cancer cells were in close contact with astrocytes, GFP+ cancer cells were plated on top of a monolayer of primary mouse astrocytes in media supplemented with 2% of CS-FBS, and green area measured over time by live imaging. Surprisingly, only H2039Br3 cells showed a significant increase in proliferation when cells were co-cultured with astrocytes, while H358, PC9Br3 and Lu65 showed similar ability to grow alone or in contact with astrocytes **(Fig 4A-D).** Nonetheless, entrectinib significantly decreased proliferation of H2030Br, Lu65 and PC9Br3 cancer cells alone or co-cultured with astrocytes (**Fig. 4B-D**). By contrast, H358 cells which were sensitive to entrectinib when stimulated with Ast-CM (**Fig 3B**), were rendered entrectinib-insensitive when cocultured with astrocytes, suggesting cell-contact with astrocytes may elicit additional survival pathways in these cells that are not targeted by entrectinib (**Fig. 4A**).

**Fig. 4.**
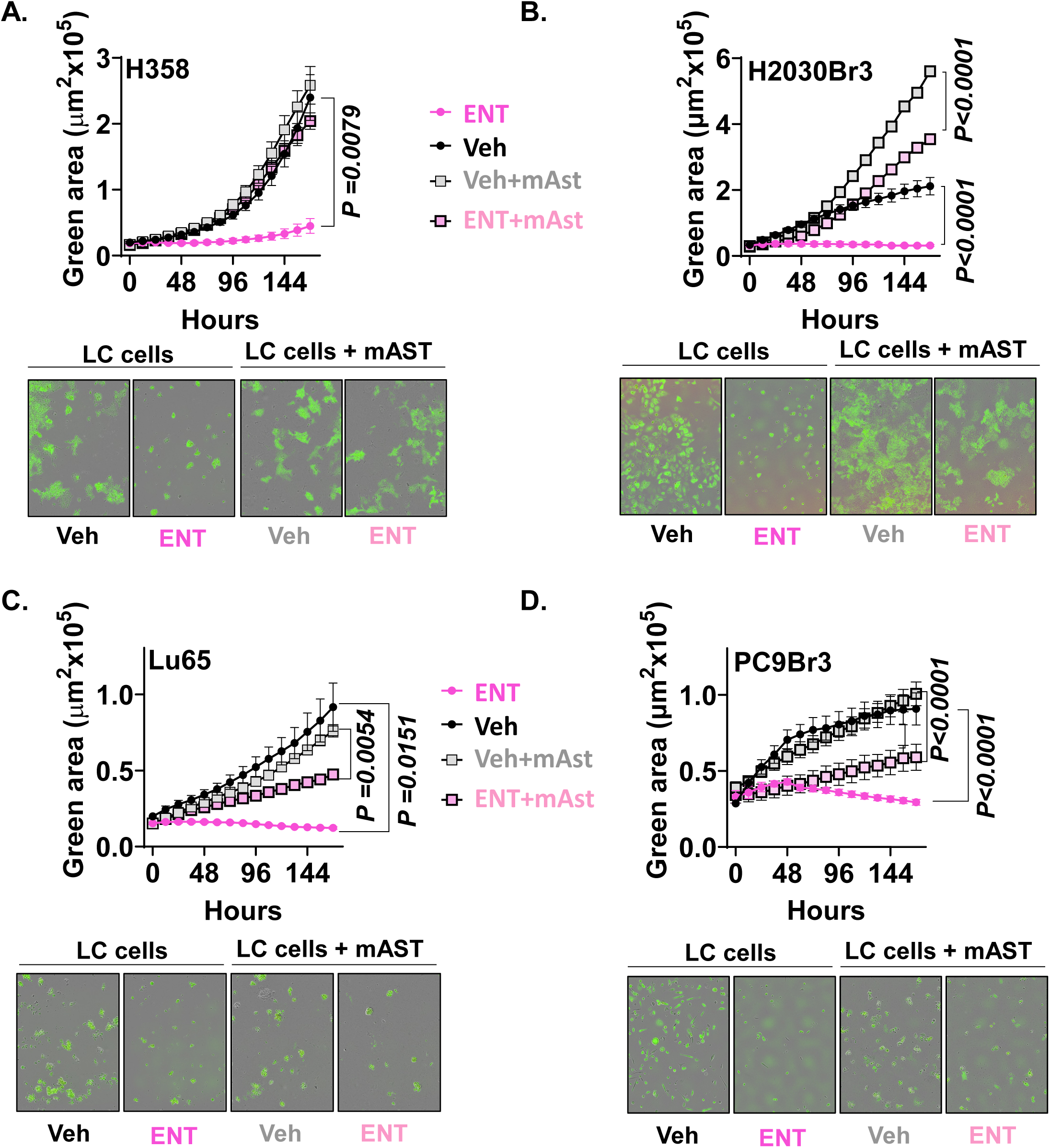
Entrectinib reduces NSCLC cell proliferation in co-culture with astrocytes. Proliferation was quantified as green area (µm²) using live-cell imaging. **(A)** H358, **(B)** H2030Br3, **(C)** Lu65, and **(D)** PC9Br3 cells were cultured either in starvation media (2% CS-FBS) alone or over a 100 % confluent monolayer of primary mouse astrocytes, in combination with vehicle (DMSO) or Entrectinib (ENT, 1 μM). Data were analyzed by repeated-measures two-way ANOVA followed by Fisher’s LSD test (n = 5 wells per treatment). P values shown are the adjusted P value for Veh vs Veh+ Ent or Veh-mAst vs ENT-mAst at the last time point.

### Entrectinib blocks brain metastatic colonization (prevents) and progression (therapeutic) of H203rM3 cells in preclinical models in vivo

We next sought to determine whether entrectinib could effectively block brain metastatic colonization (prevention of metastases) or decrease progression of existing brain metastases, using an experimental model of intracardiac injection. This model bypasses cell dissemination from the primary tumor but allows us to measure the ability of cancer cells to seed and colonize distant sites, including the brain. To determine if entrectinib could block seeding and early metastatic colonization, female NSG mice were pretreated with vehicle or 60 mg/kg BID entrectinib starting 3 days prior to intracardiac injection of 200.000 brain tropic H2030Br GFP-luc cells (which robustly form BM), and whole body IVIS was used to measure metastatic burden over time (**Fig 5A**). Entrectinib significantly decreased head metastatic progression compared to vehicle-treated mice (**Fig 5B**), which reflected a significant prevention of brain metastatic colonization as demonstrated by ex-vivo IVIS as well as histological quantification of metastases (**Fig 5C, D**). Notably, entrectinib had minimal impact on cancer cell colonization at extracranial sites with no significant differences in tumor burden in the lungs and a moderate decrease in metastatic burden in the liver-which can also produce BDNF^34^ (**Fig 5E, F**), further supporting the notion that paracrine activation of NTRK, particularly TrkB is critical to metastatic seeding and colonization in BDNF expressing organs.

**Fig. 5.**
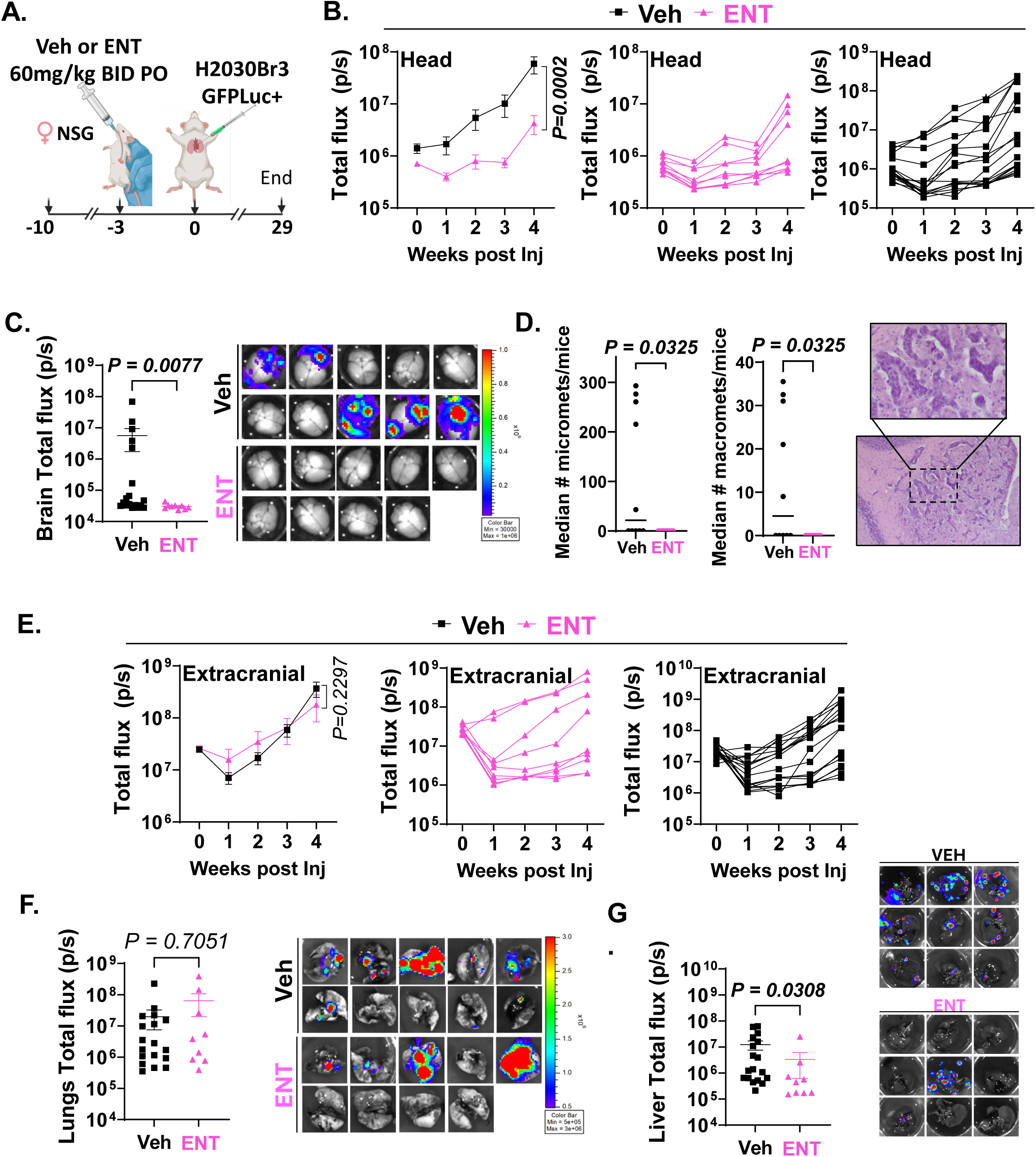
Entrectinib prevents brain metastatic colonization of H2030Br3 cells. **(A)** Experimental design: NSG female were randomized to receive vehicle or Entrectinib (ENT, 60 mg/kg, twice daily [BID], orally [PO]). Treatment began three days prior to intracardiac injection of 200,000 H2030Br3 GFPLuc cells. Metastatic progression was monitored weekly by IVIS imaging, and mice were euthanized four weeks post-inoculation. **(B)** Left graph shows mean ± SEM of total head flux by treatment; center and right graphs show individual head flux values for ENT-treated and vehicle-treated mice, respectively. **(C)** Total ex vivo flux in brain with representative IVIS images. **(D)** Each dot represents the median number of micro (<300 µm) and macro (>300 µm) metastases per mouse, Mann-Whitney test; representative H&E images of macro and micro metastases forming a cluster is shown. **(E)** Left graph shows mean ± SEM of total extracranial flux by treatment; center and right graphs show individual extracranial flux values for ENT-treated and vehicle-treated mice. Group sizes are: Vehicle = 18, ENT = 9. **(F)** Total ex vivo flux in lungs with representative IVIS images. **(G)** Total ex vivo flux in livers with representative IVIS images. Statistical analysis: In vivo IVIS data were analyzed using two-way repeated-measures ANOVA, P values show adjusted P value for the last time point; ex vivo data were analyzed using Kruskal–Wallis test followed by Fisher’s LSD post hoc test.

We next tested the effectiveness of entrectinib to decrease brain metastatic progression in the setting of well-established brain metastases. For this, H2030Br cells were injected in male and female NSG mice, brain metastases were allowed to grow for 25 days and mice with similar head tumor burden were randomized to either vehicle or entrectinib treatments (**Fig 6A**). Entrectinib decreased the progression of brain metastases as demonstrated by decreased head IVIS signal (**Fig 6B**). Similarly to the preventive setting, entrectinib did not show a significant effect on extracranial metastatic progression, further suggesting paracrine NTRK activation is critical to brain metastatic progression (**Fig 6C**). Histological analysis further shows that brains from vehicle treated mice have large and multiple panCK+ BM with high expression of p-TrkB, while entrectinib treated mice contained smaller metastases with heterogeneous but significantly lower intensity of p-TrkB (**Fig 6 D,E and Sup. Fig 3**), confirming target inhibition *in vivo*.

**Fig. 6.**
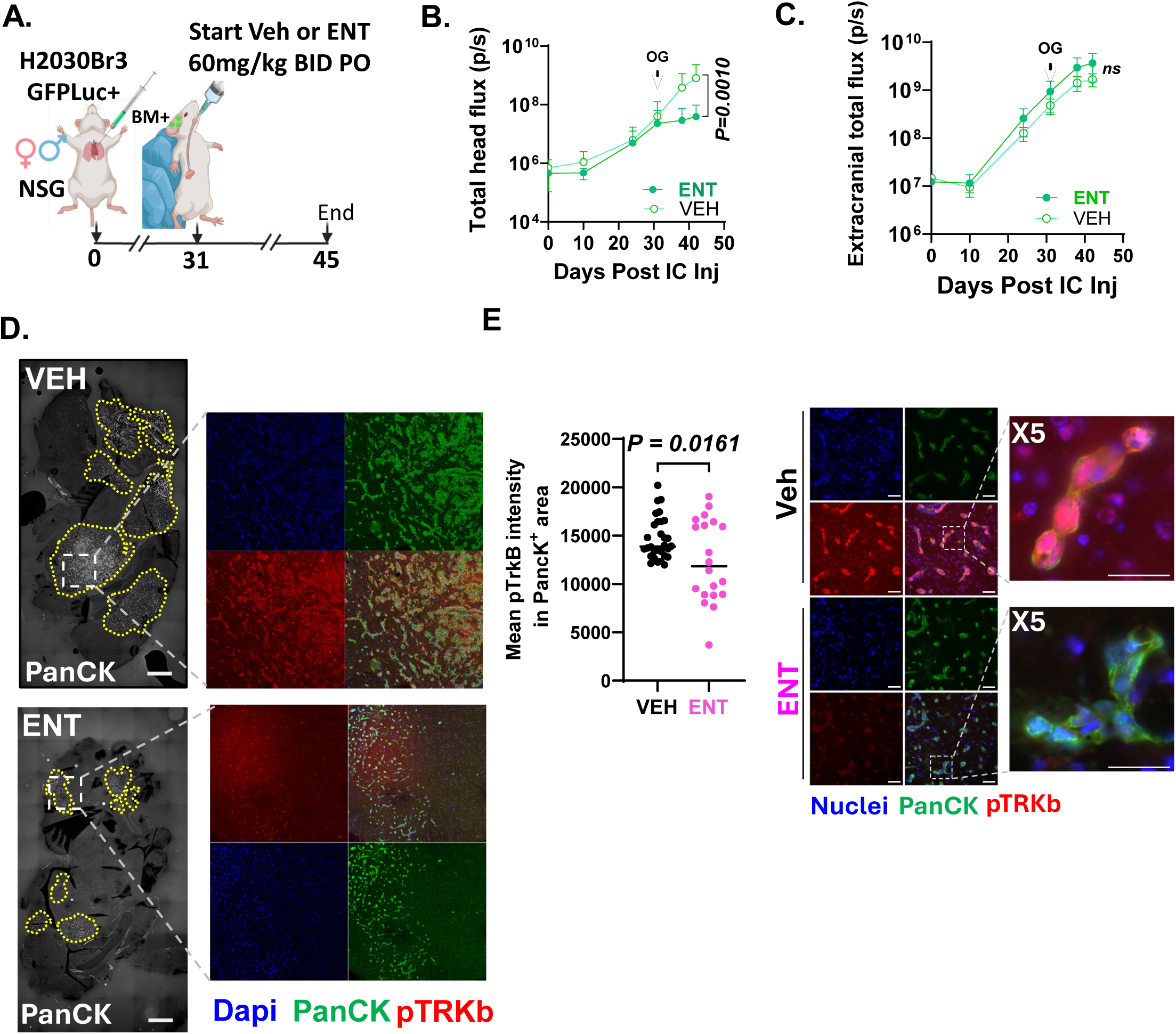
Entrectinib reduces brain metastatic burden in a late-stage model of BMs. **(A)** Experimental design: NSG female and male mice were injected intracardially with 200,000 H2030BrM3 GFPLuc cells. Head IVIS signal was monitored every 10 days. Mice showing a head signal at least twice the initial level were randomized to receive vehicle or Entrectinib (ENT, 60 mg/kg, twice daily [BID], orally [PO]). Group sizes: Female (ENT) = 6, Male (ENT) = 6, Female (Veh) = 5, Male (Veh) = 7. Treatment lasted 14 days, and metastatic burden was monitored weekly. **(B)** Total head flux for both sexes combined. **(C)** Total extracranial flux for both sexes combined. In vivo IVIS data were analyzed using two-way repeated-measures ANOVA followed by Fisher’s LSD test for multiple comparisons. P values shown are adjusted P value for last time point. **(D)** Representative images of full brain sagittal sections from vehicle and entrectinib treated mice. Dotted lines mark the pan-cytokeratin positive (PanCK+, gray) tumor areas. Boxes show 5X magnification of p-TrkB (red), panCK (green), dapi (nuclei) and merged images. Scale bar=1000 um **(E).** Quantification of p-TrkB in 10 images/mice from 2 individual mice per group, at 40X. PanCK+ mask was generated to define tumor areas and converted into a region of interest (ROI), pTrkB mean fluorescence intensity (MFI) was measured within pan-CK+ areas only. Representative images of fields used for quantification. Scale bar = 50 um

## DISCUSSION

While the tumor promoting role for NTRK receptors in lung cancer progression is well recognized, our studies further demonstrate a unique role for brain activity of NTRK in lung cancer, and the potential use of FDA-approved NTRK inhibitors in the prevention of CNS metastases for non-NTRK rearranged tumors. The fact that NTRK expression can be detected in primary tumors and does not significantly change in their brain metastases suggests that testing for WT NTRK expression could serve as a criterion to define patients who could benefit from preventive strategies with NTRK inhibitors.

Mechanistically, we demonstrate that entrectinib can block growth of lung cancer cells in the presence of astrocytes, cells that provide multiple survival signals and are important components of the brain niche^35,36^. This agrees with prior findings in preclinical models of breast cancer brain metastases showing that astrocytes activate TrkB and also other receptors in cancer cells (i.e EGFR) promoting proliferation, invasion and brain colonization in breast cancer^7,31^. The observation that entrectinib is highly effective in blocking pro-survival signaling and cancer cell proliferation in multiple cell lines, when cells are cultured alone, in comparison to cells in co-culture with astrocytes, highlights the important contribution from cells in the tumor microenvironment to the activation of growth and survival signals, and the need to consider how tumor-host interactions alter therapeutic responses in brain metastases. The significant effect of entrectinib blocking brain (the organ with highest levels of BDNF) and to a lesser extent liver metastases (which express low levels of BDNF) without altering lung metastases, highlights the potential to exploit the paracrine activation of TKIs in an organ specific manner. Specifically, these data suggest NTRK inhibitors may have a role in protecting the brain (and potentially the liver) from metastatic spread, which could and should be explored clinically.

While our data suggest a role for microenvironment activation of NTRK to support growth, it is likely that additional mechanisms contribute to the pro-metastatic function of NTRK in brain metastases. For example, recent research indicates that primary brain tumors exploit BDNF-TrkB signaling to modulate cancer–neuron synapses, and inhibiting this pathway significantly suppresses tumor growth^37,38^. Given that neuron-to-lung and neuron-to-breast cancer synapses are essential for brain metastatic progression^39,40^, it is plausible that BDNF/TrkB-mediated synaptic plasticity and connectivity also play a role in the initiation of CNS metastases. In fact, the effective abolishment of brain metastases when entrectinib is used prior to tumor injection suggests these early cancer-neuron or cancer-astrocyte interactions may be key for NTRK-promotion of brain colonization. Nonetheless, our findings that entrectinib also delayed the progression of late-stage metastases suggest NTRK activation remains active throughout metastatic progression and can be targeted in a preventive and therapeutic setting.

Our results showed that entrectinib was most effective than the most selective TrkB inhibitors (such as cyclotraxin and ANA-12) in inhibiting proliferation of lung cancer cells. It is likely than entrectinib a ATP□competitive tyrosine kinase inhibitor with favorable intracellular bioavailability, is able to achieve sustained inhibition of the Trk kinase domain, while ANA□12 (which interferes with ligand binding^41^), and cyclotraxin□B (which acts as a negative allosteric modulator^42^); are non□ATP□competitive and may produce only partial or transient suppression of TrkB signaling. Moreover, while we focused on TrkB function here due to its increased expression among lung cancers and cell lines, it is possible that TrkA also plays a role in the sensitivity to panTrk inhibitors. Elevated TrkA protein expression and increased NGF levels have been observed in squamous cell carcinoma compared to benign lesions and other malignant lung cancer histological subtypes^43^. It is therefore possible that even low□abundance TrkA, when stimulated, could contribute to residual AKT/ERK signaling when TrkB is selectively inhibited. Furthermore, recent studies indicate that suppression of wild-type NTRK1 in murine lung cancer models enhances sensitivity to immunotherapy by promoting complement C3-mediated activation of T cells^44^. Since our studies were limited to human xenograft studies in immune-compromised mice, the extent to which entrectinib impacts brain metastatic progression and its impact on anti-tumoral responses remain unknown.

Finally, among the tested NTRK inhibitors in clinical development, entrectinib blocked NTRK signaling at a lower effective dose, consistent with prior reports of increased IC50 for this compound. However, it is possible that higher effective doses of other TKIs have similar ability to entrectinib to target brain metastatic colonization. Additional studies are needed to define whether LOXO-101 or new generation NTRKs at their effective doses *in vivo*, have similar ability to prevent the seeding and colonization and decrease progression of existing brain metastases. While the side effects for targeting the normal NTRK function in the brain including pain sensation, appetite regulation, proprioception, memory and learning need to be considered, the potential value for NTRK inhibition for the prevention of brain metastases may open new opportunities for prevention of BM in patients at high risk.

## Authorship contribution

Study supervision: DC; Conception and design: DC, MJC; Data acquisition, analysis, resources: DC, MJC, JAJ, RAM, TP, SK, DRO, ACN, RAN, DRC. All authors contributed to writing and approved the manuscript.

## Acknowledgements

We thank Dr. Sharon Pine for sharing the H2030Br3 and PC9Br3 cells. We thank Dr. A. Van Bokhoven at the Biorepository Core Facility and personnel at the University of Colorado Neurosurgery Nervous System Biorepository for providing de-identified human tissue. We thank the University of Colorado Cancer Center Animal Imaging shared resources supported by NCI P30CA046934 and CTSA UL1TR001082 Center grants.

