## Supplementary Table 1 for "Targeting wild type NTRK decreases brain metastases of lung cancers non-driven by NTRK fusions"

**Supplementary Table 1: List of antibodies, sources, and working dilutions used in this study**

| Target | Vendor | Cat# | Dilution/Use | RRID |
| --- | --- | --- | --- | --- |
| pTrkB (Y515) | Invitrogen | PA5-36695 | 1:1000/WB  1:150/IF | AB_2553666 |
| pTrkB (Y816) | Millipore | ABN1381 | 1:1000/WB | AB_2721199 |
| TrkB | ProteinTech | 13129-1-AP | 1:1000/WB  1:8000/IHC | AB_2155156 |
| TrkA | Abcam | ab76291 | 1:1000/WB | AB_1524514 |
| TrkC | Cell Signaling | 3376 | 1:1000/WB | AB_2155283 |
| ROS1 | Cell Signaling | 3287 | 1:1000/WB | AB_2797603 |
| ALK | Cell Signaling | 3791 | 1:1000/WB | AB_1950402 |
| AKT | Cell Signaling | 9272S | 1:1000/WB | AB_329827 |
| pAKT(S473) | Cell Signaling | 4060S | 1:1000/WB | AB_2315049 |
| ERK | Cell Signaling | 9102 | 1:1000/WB | AB_330744 |
| pERK(T202/T204) | Cell Signaling | 9101 | 1:1000/WB | AB_331646 |
| Tubulin | Sigma | T5168 | 1:10000/WB | AB_477579 |
| Goat anti-Mouse Alexa Fluor 680 | Life Technology | A21058 | 1:10000/WB | AB_2535724 |
| Goat anti-Rabbit Alexa Fluor 680 | Life Technology | A21109 | 1:10000/WB | AB_2535758 |
| Goat anti-Mouse Alexa Fluor 800 | LICOR | 926-32210 | 1:10000/WB | AB_621842 |
| Cytokeratin | Dako | M0821 | 1:200/IF | AB_2858276 |
| GFAP | Invitrogen | 13-0300 | 1:500/IF | AB_86543 |
| BDNF | Abcam | Ab108319 | 1:100/IF | AB_10862052 |
| Donkey anti rabbit Alexa Fluor 555 | Jackson ImmunoResearch Labs | 711-565-152 | 1:500/IF | AB_3095471 |
| Donkey anti mouse Alexa Fluor 480 | Jackson ImmunoResearch Labs | 715-545-151 | 1:500/IF | AB_2341099 |
| Donkey anti rat Alexa Fluor 480 | Jackson ImmunoResearch Labs | 712-545-153 | 1:500/IF | AB_2340684 |
