## Supplementary Figure 1 for "Targeting wild type NTRK decreases brain metastases of lung cancers non-driven by NTRK fusions"

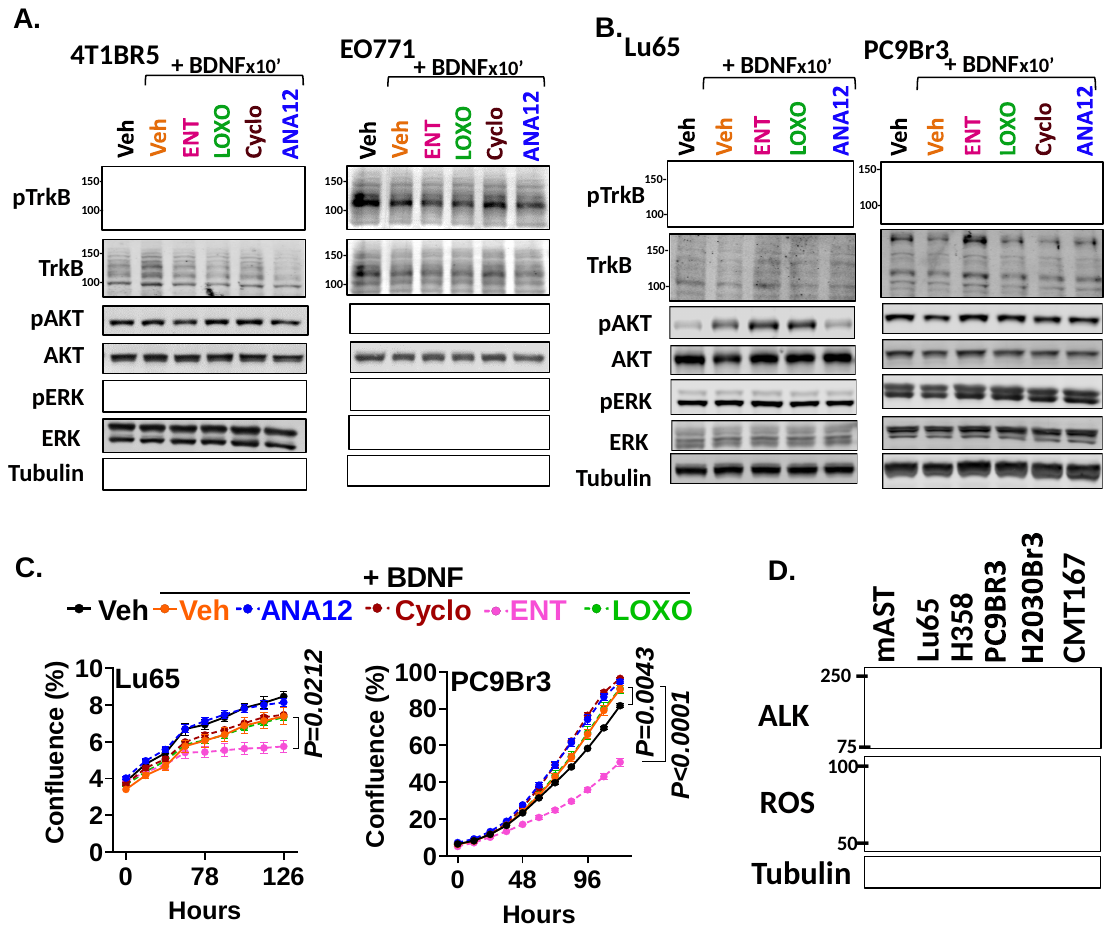

**Sup. Fig 1.** **ENT decreased proliferation independently of AKT or Erk signaling pathway in some NSCLC.**

Western blot of serum-starved murine triple negative breast cancer cells 4T1Br5 and E0771 **(A)** and NSCLC Lu65 and PC9Br3 cells **(B)**  pretreated for 2 hours with DMSO (Veh), Entrectinib (ENT), LOXO101 (LOXO), Cyclotraxin (Cyclo), or ANA12 (all drugs at 1 μM), followed by stimulation with 50 ng/mL BDNF for 10 minutes. Tubulin was used as a loading control in all western blots. **C.** Graphs show proliferation as the media percentage of confluency ± SEM measured using Incucyte live-cell imaging (n = 5 per treatment) for cells cultured in starvation media supplemented with 2% CS-FBS and treated as in (B). Data were analyzed using repeated-measures two-way ANOVA followed by post hoc multiple-comparison corrections. **D.** Western blot of ALK and ROS in NSCLC cell lines.
