## Supplementary Figure 2 for "Targeting wild type NTRK decreases brain metastases of lung cancers non-driven by NTRK fusions"

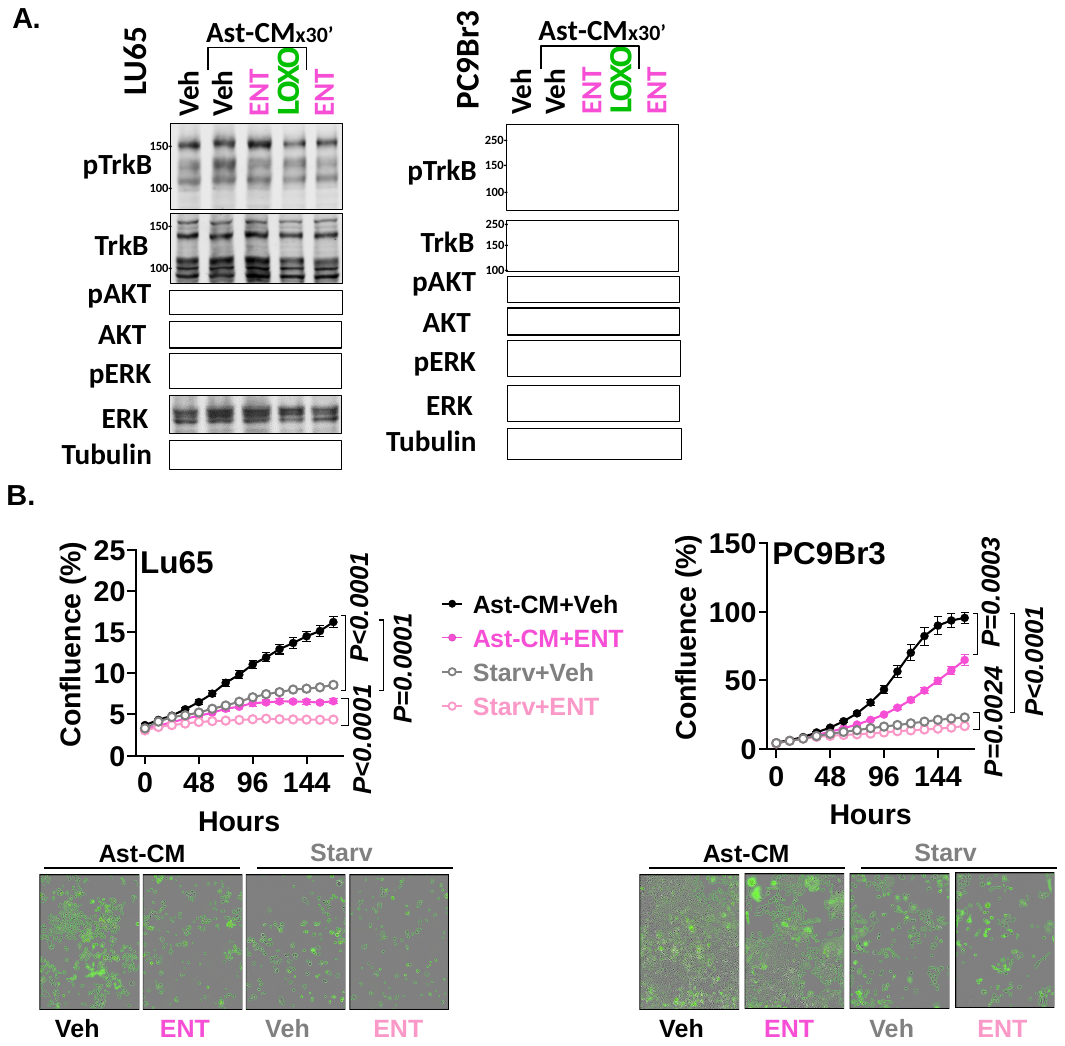

**Sup. Fig 2**

**A.** TrkB signaling in NSCLC cells serum-starved overnight, pretreated for 2 hours with 1 μM Entrectinib (ENT) or LOXO101 (LOXO), followed by stimulation with Ast-CM (10X) for 30 minutes. Tubulin was used as a loading control. **B.** Proliferation expressed as percentage of confluence in NSCLC cells cultured in serum-starved media and treated or not with Ast-CM (10X) in combination with vehicle (DMSO) or ENT at 1 μM. Data were analyzed using repeated-measures two-way ANOVA followed by Fisher’s LSD test (n = 5 wells per treatment).
