## Supplementary Figure 3 for "Targeting wild type NTRK decreases brain metastases of lung cancers non-driven by NTRK fusions"

**Sup. Fig 3. Workflow for quantifying pTRKB fluorescence intensity within GFP-defined ROIs.**

Representative images illustrating the image analysis pipeline used for fluorescence quantification in ImageJ. The GFP channel (top left), marking PanCK-positive brain mets, was used to generate a segmentation mask. After manual threshold adjustment, the GFP image was converted into a binary mask (top right) to identify GFP-positive regions. The refined mask was then converted into regions of interest (ROIs) and overlaid onto the RFP channel image representing pTRKB immunofluorescence (bottom left). Finally, the ROIs were applied to the RFP image to quantify mean fluorescence intensity and area of pTRKB signal specifically within GFP-positive regions (bottom right, overlay showing ROIs). Measurements from all analyzed images were automatically exported and compiled into a CSV file for downstream statistical analysis.
